# Functional Separation of mRNA Domains Coordinates Pluripotent Cell Behavior

**DOI:** 10.1101/2022.03.15.484504

**Authors:** Ze Yang, Shaoyi Ji, Kristina Ivanov, Priyanka Kadav, Mengmeng Song, Leonardi Gozali, Sophie Parsa, Barry Behr, Mary A. Hynes

**Affiliations:** Stanford University Department of Biology and the Stanford IVF laboratory, Stanford University, Stanford California 94305; Department of Obstetrics and Gynecology and the Stanford IVF laboratory, Stanford University, Stanford California 94305

## Abstract

Differential expression of mRNA coding sequences (CDSs) and 3’untranslated regions (UTRs) is widespread, yet whether these domains contribute independently to cellular function remains unclear. Here, using *Nanog* as a model, we find that *Nanog* mRNA domain usage is spatially organized in mouse and human pluripotent cells and in the blastocyst, with cells enriched in 3’UTR transcripts marking colony borders and cells enriched for the *Nanog* CDS located interior. Functional perturbation reveals a marked asymmetry between mRNA domains. Loss of the *Nanog* 3’UTR leads to defects in colony architecture, cell spreading and morphogenetic behavior, accompanied by decreased extracellular matrix modeling gene expression and ROCK dependent cytoskeletal organization. In contrast, deletion of the *Nanog* CDS primarily disrupts epithelial polarity-associated transcriptional programs and the expression of chromatin regulators, consistent with a dominant role for the Nanog protein in transcriptional and epigenetic control. These domain specific effects are not redundant but instead reflect separable regulatory activities encoded within a single transcript. Together, these findings demonstrate that distinct regions of a single mRNA can encode separable and asymmetric biological functions, revealing mRNA domain usage as a distinct regulatory layer through which genes can encode multiple biological outputs beyond protein coding capacity.

## Introduction

Messenger RNAs (mRNAs) contain both coding sequences (CDS) and untranslated regions (UTRs), with biological function most often ascribed to the encoded protein. However, we and others have shown that 3’UTRs and their cognate CDSs can be differentially expressed across tissues and developmental contexts, forming striking and often reciprocal spatial pattens of expression (Mercer, Wilhelm et al. 2011, Kocabas, Duarte et al. 2015, Malka, Steiman-Shimony et al. 2017, Ji, Yang et al. 2021). As a result, many cells harbor abundant, stable, mRNA derived noncoding 3’UTR fragments (Malka, Alkan et al. 2022), raising the possibility that such 3’UTR RNA components may carry biological functions beyond regulating transcript stability or localization.

Isolated 3’UTR components can be generated through cleavage and 5’ recapping mechanisms (Nejc Haberman 2023), whereas isolated CDS components, often with shortened 3’UTRs, can arise through alternative polyadenylation pathways (Mayr 2017). Despite the prevalence of differential mRNA component expression, whether isolated 3’UTRs perform distinct biological functions independent of their cognate coding regions remains largely unresolved.

Here, we focus on the core pluripotency genes *Sox2*, *Oct4*, and *Nanog* (the SON genes) which together sustain embryonic stem cell (ESC) identity and developmental potential (Chambers and Tomlinson 2009, Festuccia, Osorno et al. 2013, Cancedda and Mastrogiacomo 2025). *Sox2* and *Oct4* are essential for pluripotency both *in vivo* and *in vitro*, with deviations in their precisely tuned expression levels leading to differentiation (Niwa, Miyazaki et al. 2000, Loh, Wu et al. 2006). Nanog, although long considered a core pluripotency factor (Loh, Wu et al. 2006, Cancedda and Mastrogiacomo 2025), plays a more nuanced role. It is dispensable for ESC self-renewal under standard in vitro conditions (Hatano, Tada et al. 2005, Chambers, Silva et al. 2007), yet it is essential *in vivo*, where blastocysts fail in its absence (Rossant and Tam 2009) and during induced pluripotent stem (iPS) cell reprogramming (Mitsui, Tokuzawa et al. 2003, Chambers, Silva et al. 2007, Silva, Nichols et al. 2009). Despite extensive molecular characterization of SON protein interactions (Cancedda and Mastrogiacomo 2025), the specific developmental function of Nanog has remained incompletely defined.

We find that among the SON genes, *Nanog* shows the most pronounced differential usage of its 3’UTR and CDS components in both mouse and human ESCs. This pattern occurs under standard culture conditions, in single-cell RNA-seq datasets, and in the blastocyst, and is further enhanced when ESCs are organized into two-dimensional (2D) colonies. Specifically, cells at colony border cells express high levels of the *Nanog* 3’UTR with minimal detectable CDS, while adjacent interior cells display the opposite pattern.

Functional perturbation studies reveal that selective deletion of the *Nanog* 3’UTR downregulates extracellular matrix (ECM) remodeling programs, resulting in compact, poorly spreading colonies with disrupted actin and adhesion-associated protein localization. Overexpression of the *Nanog* 3’UTR enhances border localization, while ROCK kinase inhibition rescues these phenotypes, placing the *Nanog* 3’UTR upstream of cytoskeletal and adhesion control pathways. In contrast, deletion of the *Nanog* CDS preferentially alters epithelial polarity and chromatin associated programs, consistent with the canonical role of Nanog protein in restraining epithelial reorganization within pluripotent colonies.

Together, these findings demonstrate that differential mRNA component expression at the *Nanog* locus encodes distinct and separable regulatory function, revealing a previously unrecognized role for a 3’UTR-derived RNA in controlling embryonic stem cell morphogenetic behavior.

## Results

### *Nanog* mRNA domains exhibit spatially organized expression in pluripotent colonies

To determine whether differential expression of mRNA coding sequences (CDSs) and 3’untranslated regions (3’UTRs) contributes to pluripotent stem cell organization, we analyzed expression patterns of the core pluripotency genes *Nanog*, *Sox2,* and *Oct4* in mouse and human embryonic stem cells (ESCs). Among these genes, *Nanog* exhibited the most pronounced and reproducible segregation of its 3’UTR and CDS components across cells. We therefore focused subsequent analyses on *Nanog* to assess whether differential mRNA domain expression at this locus has distinct functional consequences.

SON gene 3’UTR and CDS mRNA component expression was examined by dual color in situ hybridization (ISH) in mouse and human ESCs, with CDS detected in green and 3’UTR detected in red. Cells were cultured under standard planar culture conditions or in two dimensional circular micropatterns to enhance spatial organization (CYTOO™ platform) (Warmflash, Sorre et al. 2014, Deglincerti, Croft et al. 2016, Morgani, Metzger et al. 2018, Ji, Yang et al. 2021). Under micropatterned conditions, *Nanog* exhibited striking spatial segregation of its mRNA domains, with high 3’UTR cells enriched at colony borders and high CDS cells occupying interior positions (Figure 1A-C). In contrast *Oct4* and *Sox2* displayed weaker and less consistent spatial organization (Figure1-figure supplement 1A).

**Figure 1.**
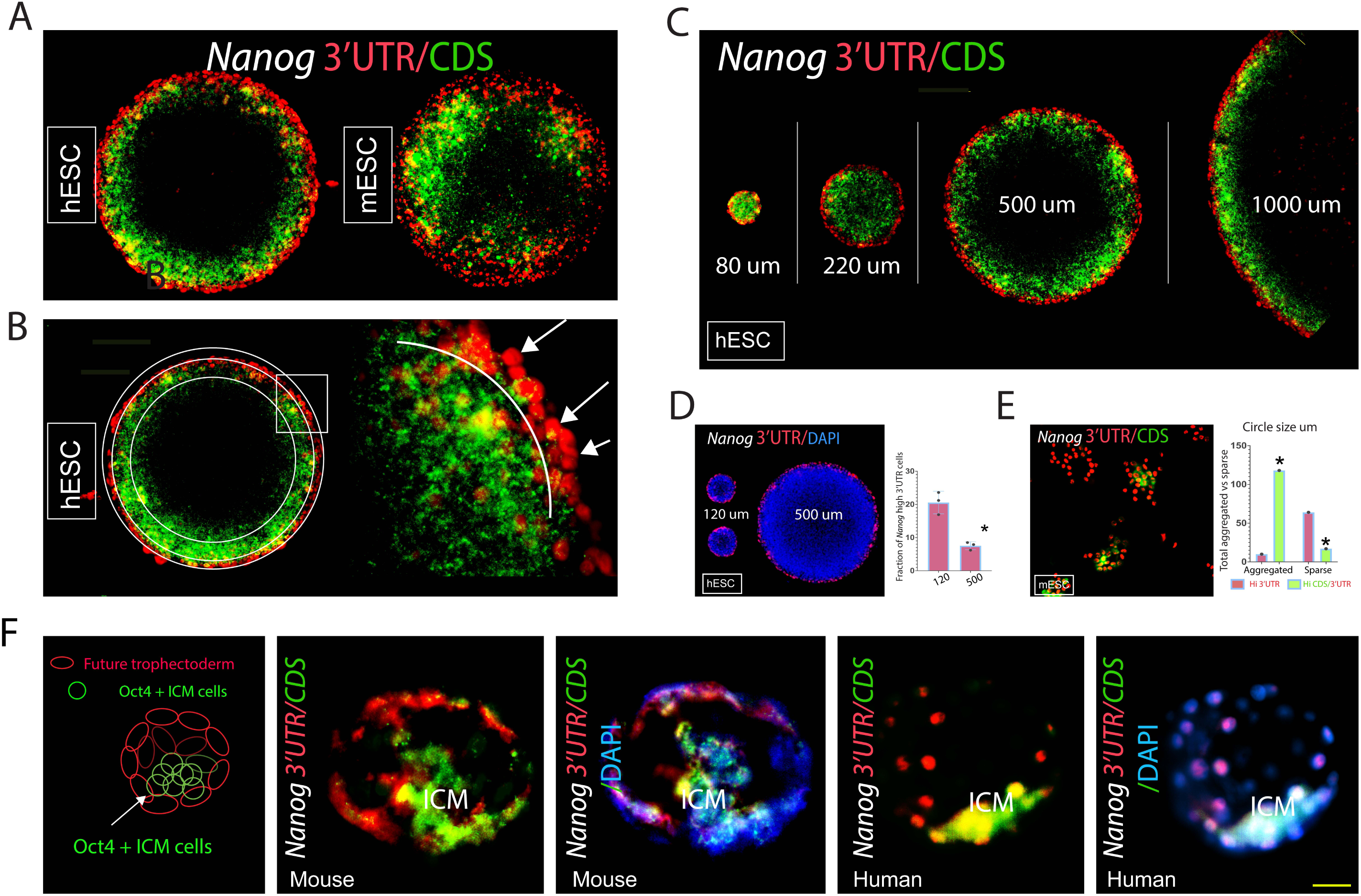
Spatial segregation of *Nanog* mRNA domains in pluripotent colonies and the blastocyst. Fluorescence in situ hybridization (ISH) for *Nanog* using 3’UTR specific (red) and CDS specific (green) probes in mouse and human embryonic stem cells (ESCs) and blastocyst stage embryos. (A) Representative images showing spatial distribution of *Nanog* 3’UTR and CDS expression patterned ESC colonies. *Nanog* exhibits prominent most pronounced spatial segregation of 3’UTR and CDS biased expression compared with other pluripotency associated genes. (See Figure 1-figure supplement 1A for *Sox2* and *Oct4*). (B) Higher magnification image of *Nanog* expression at colony border highlighting cells with high 3’UTR signal and little or no detectable CDS signal (arrows). (C) *Nanog* 3’UTR/CDS spatial patterning is maintained across colony diameters ranging from 80-1000 um (images from different micropatterned chips). (D) Quantification of the fraction of high *Nanog* 3’UTR cells as a function of colony size after plating of a single pool of hESCs (mean +/− SEM; n=3 biological replicates; Student’s t-test). (E) *Nanog* 3’UTR/CDS expression patterns differ between aggregated (>3 cells) and sparsely distributed ESCs (means of three biological replicates: Chi-square analysis (X^2^, p<0.01)). (F) *Nanog* 3’UTR/CDS expression in blastocyst stage mouse and human embryos. Additional markers are shown in Figure 1-figure supplement 1A. Scale bar: A, C; 100 um, B; left 100 um, right 20 um, C; 160 um, E; 130 um, F; mouse 20 um, human, 45 um.

*Nanog* 3’UTR and CDS spatial patterning was highly robust across hundreds of micropatterned colonies and across a wide range of colony diameters (Figure 1C; Figure1-figure supplement 2D,E). Notably, the combined width of the *Nanog* expression domain (3’UTR plus CDS) remained approximately 100 um across colony sizes. In smaller colonies, *Nanog* expression filled the colony, whereas in colonies larger than 500 um, central cells showed low or undetectable SON mRNA component expression (Figure1A-C, Figure 1-figure supplement 1A), consistent with edge-initiated patterning in pluripotent colonies (Warmflash, Sorre et al. 2014).

To determine whether *Nanog* mRNA domain usage reflects intrinsic cell states or responds to local cellular context, a single pool of ESCs was plated onto micropatterned chips containing circles of various diameters. Despite being derived from the same initial population, smaller colonies contained a significantly higher fraction of high *Nanog* 3’UTR cells relative to total cell number (Figure 1D). Analysis of colonies with variable local density (arising occasionally from uneven plating) further revealed that high *Nanog* 3’UTR expression was observed under conditions in which interior high *Nanog* CDS expression was low or absent (Figure 1-figure supplement 2B,C,E), consistent with modulation of mRNA domain usage by local environment.

Similar context dependent patterns of *Nanog* mRNA domain expression were observed across multiple biological settings. Under standard culture conditions dispersed ESCs preferentially expressed high *Nanog* 3’UTR, whereas aggregated cells were significantly more likely to expression high *Nanog* CDS (Figure 1E). Likewise, in both mouse and human blastocysts, inner cell mass (ICM) cells expressed high *Nanog* CDS (along with some 3’UTR), whereas outer cells predominantly expressed the 3’UTR (Figure 1F, Figure 1-figure supplement 1B-D). Consistent patterns were also evident in single-cell RNA sequencing datasets (Figure 2A-D, Figure 2-figure supplement 1A-C). Across all contexts examined, cells surrounded by neighboring cells preferentially expressed the *Nanog* CDS, whereas isolated or border cells predominantly expressed the *Nanog* 3’UTR, indicating that *Nanog* mRNA domain usage varies systematically with local cellular context.

**Figure 2.**
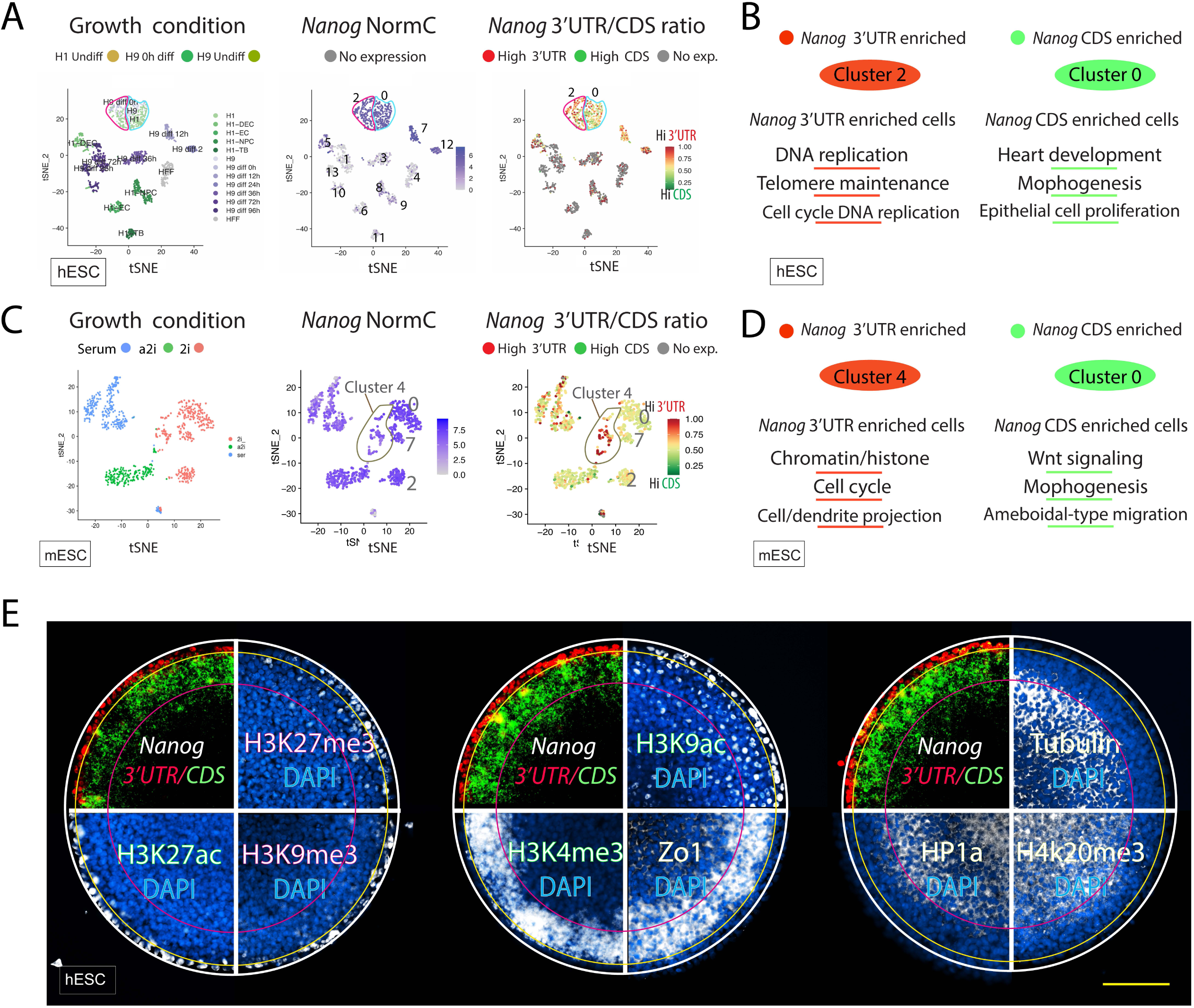
Differential *Nanog* mRNA domain usage stratifies pluripotent cell states. (A) t-SNE projections of hESCs cultured under indicated growth conditions, colored by growth condition (left), normalized *Nanog* expression (middle), and relative *Nanog* mRNA domain usage (3’UTR vs CDS; right). Red indicates cells with higher relative 3’UTR usage and green indicates cells with higher relative CDS usage. (B) Gene Ontology (GO) terms associated with hESC clusters characterized by higher relative *Nanog* 3’UTR or CDS usage. (C) t-SNE projections of mESCs under indicated growth conditions, colored as in (A). (D) GO terms associated with mESC clusters characterized by higher relative *Nanog* 3’UTR or CDS usage. (E) Combined ISH and immunofluorescence showing spatial organization of *Nanog* mRNA domain usage together with indicated chromatin and junctional markers. IF signals align with *Nanog* 3’UTR/CDS expression in concentric regions. Additional IF analyses are shown in Figure 2-figure supplement 1D-H). Scale bar E; 100 μm.

### *Nanog* 3’UTR enriched cells exhibit distinct transcriptional profiles in mouse and human ESCs

To examine transcriptional differences associated with *Nanog* mRNA domain usage, we analyzed publicly available mouse (Kolodziejczyk, Kim et al. 2015) and human (Chu, Leng et al. 2016) single cell RNA-sequencing (scRNA-seq) datasets generated using full-length transcript coverage (Smart-seq). We developed a pipeline that overlays gene-specific 3’UTR/CDS expression ratios onto normalized counts (NormC)-based clustering (Figure 2B,D; Figure 2-supplement figure 1B,C; see Methods). First, transcriptome wide analysis (mESCs) shows no overall correlation between NormC gene expression and 3’UTR/CDS ratio (Figure 2-figure supplement 1A). Many genes show differential 3’UTR and CDS usage in pluripotent ESCs and the ratio lookup outputs for all expressed genes in these mESC and hESC datasets are provided (<u>here</u> and <u>here</u>).

Overlay of the *Nanog* 3’UTR/CDS ratio onto NormC-clustered cells revealed discrete populations enriched for *Nanog* 3’UTR expression in both human (cluster 2) and mouse (cluster 4) ESCs (Figure 2B,D), with these clusters classified as undifferentiated. For mouse ESCs, subsequent analyses focused on clusters grown under conditions most comparable to those used in the present study (clusters 0,2,4,7). Notably, overall *Nanog* mRNA abundance (NormC) did not vary substantially across these clusters (Figure 2C; middle), whereas *Nanog* mRNA domain usages differed markedly: cells in cluster 4 showed enriched *Nanog* 3’UTR expression (red), while cells in clusters 0,2,7 expressed higher levels of *Nanog* CDS (Figure 2C; right).

Differential gene expression comparing high *Nanog* 3’UTR and high *Nanog* CDS cells (hESC cluster 2 vs 0; mESC cluster 4 vs 0) identified distinct transcriptional programs shared across species. High *Nanog* 3’UTR cells showed elevated expression for genes involved in chromatin organization, cell cycle regulation, and cellular projections (Figure 2 B,D, Figure 2 supplement 1B,C). In contrast high *Nanog* CDS cells showed elevated expression of genes involved in morphogenesis, WNT signaling, and epithelial proliferation (Figure 2 B,D, Figure 2-figure supplement 1B,C). Thus, ESCs enriched for *Nanog* 3’UTR or CDS occupy transcriptionally distinct, yet overlapping, regions of pluripotent state space.

### Chromatin mark expression differs between *Nanog* mRNA domain-associated cell populations

Chromatin state is closely linked to pluripotency and differentiation, and chromatin marks can vary across cells with distinct transcriptional configurations (Vogelauer, Wu et al. 2000, Seligson, Horvath et al. 2005, Kurdistani 2011). In light of the transcriptional differences observed between *Nanog* 3’UTR and CDS enriched cells (Figure 2B,D), we next examined chromatin mark expression in ESCS cultured on micropatterned substrates. Larger colonies are shown to illustrate the spatial correspondence between *Nanog* mRNA domain usage and chromatin mark distribution.

*Nanog* 3’UTR enriched border cells and more interior *Nanog* CDS enriched cells displayed distinct chromatin mark distributions. *Nanog* CDS enriched cells preferentially expressed histone modifications associated with transcriptionally accessible chromatin (active), including robust H3K4me3 (Figure 2E; middle, Figure 2-figure supplement 1D-H). In contrast, *Nanog* 3’UTR enriched border cells expressed a combination of active (H3K9ac) and repressive (H3K9me3) histone marks, consistent with bivalent chromatin configurations characteristic of pluripotent cells (Bernstein, Mikkelsen et al. 2006) (Figure 2E, left; Figure 2-figure supplement 1D-H).

In smaller colonies, comparable in size to the in vivo blastocyst, *Nanog* mRNA component expression was observed throughout the colony (Figure 1C; Figure 1-figure supplement 2D,E), however, more centrally located cells in large colonies exhibited reduced expression of *Nanog* mRNA components and also show greatly diminished expression of chromatin marks associated with *Nanog* expressing regions. Instead, these interior cells express HP1α, H4K20me3, and βIII Tubulin, markers previously associated with transition from pluripotency and early differentiation (Figure 2E; right, Figure 2-figure supplement 1I-K).

### *Nanog* 3’UTR and CDS knockout lines exhibit distinct functional behaviors

To test the functional contributions of the *Nanog* mRNA domains, we generated selective CRISPR deletions targeting either the *Nanog* CDS or the *Nanog* 3’UTR in mouse embryonic stem cells (mESCs). mESCs were used because these cells express a single *Nanog* genotype, whereas hESCs express multiple *Nanog* pseudogenes that complicate domain specific targeting.

For *Nanog* 3’UTR deletion, two fluorescently labeled guide RNAs flanking the 3’UTR (while sparing the polyadenylation signal) were introduced, followed by FACS enrichment. *Nanog* CDS deletion was achieved using a single guide RNA (Figure 3A; left). Independent isogenic lines were established for each genotype: control (Ctl; C2, C5), *Nanog* 3’UTR knockout (NUKO; NU3, NU4), and *Nanog* CDS knockout (NCKO; NC3, NC31).

**Figure 3.**
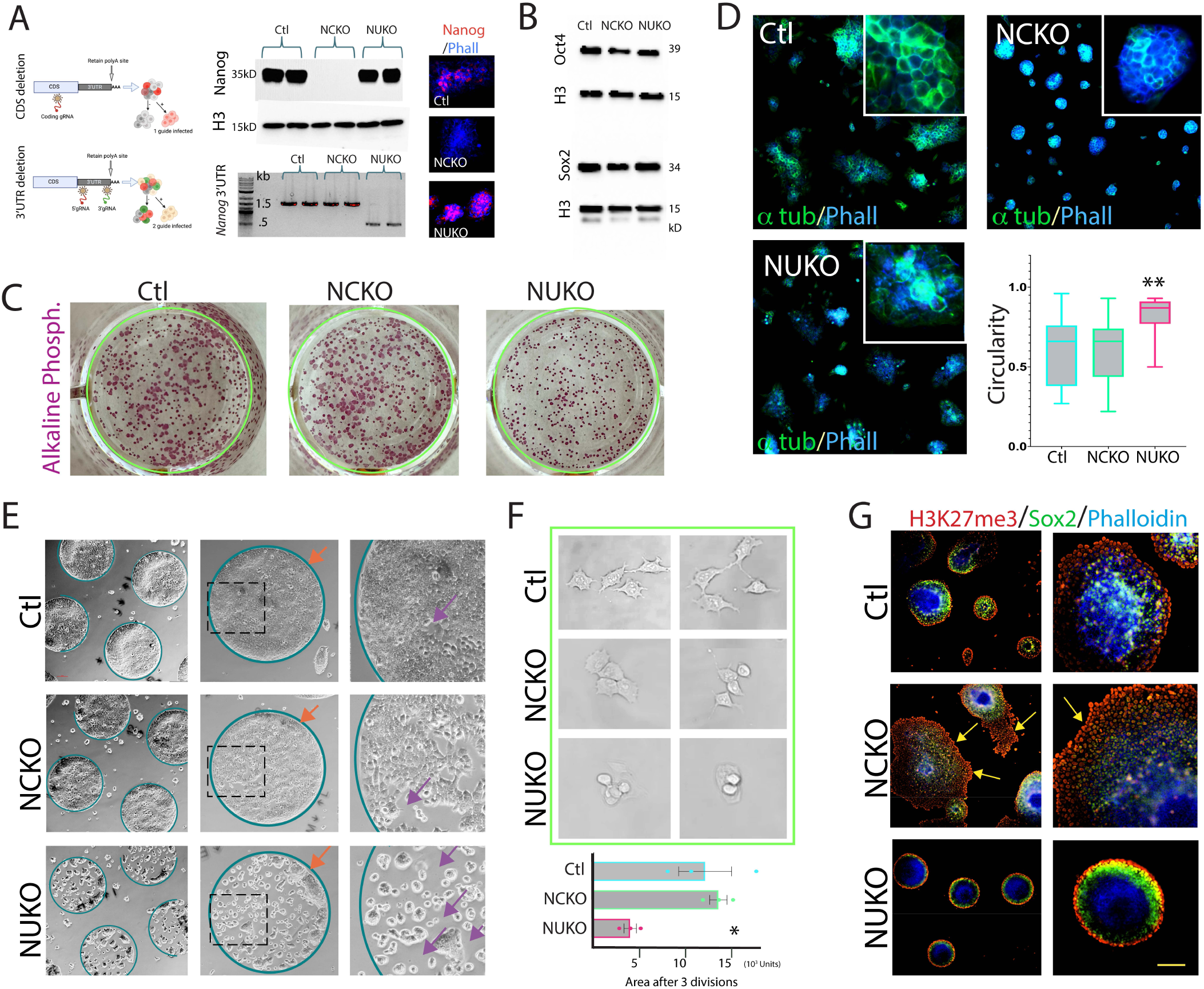
*Nanog* 3’UTR deletion impairs ESC colony dispersal. (A) Schematic of selective CRISPR knockout of the *Nanog* CDS or *Nanog* 3’UTR in mESCs with validation by genomic PCR, western blot, and *Nanog* IF. The CDS was targeted with a single gRNA and the *Nanog* 3’UTR (excluding the polyadenylation site) was deleted using two flanking gRNAs. (B) Western blot analysis of Oct4 and Sox2 expression across control (Ctl), *Nanog* CDS knockout (NCKO), and *Nanog* 3’UTR knockout (NUKO) lines indicating maintenance of pluripotency associated protein expression. (C) Low-density clonal cultures stained for alkaline phosphatase demonstrating pluripotent identity across lines. (D) IF for phalloidin and α-tubulin in planar culture. NUKO colonies show increased circularity compared to Ctl and NCKO colonies. Data are shown as mean +/− SEM (n=3 biological replicates; n>5 colonies per condition; Student’s t-test). (E) On CYTOO™ micropatterned circles, NUKO cells showed reduced spreading across the circle compared to Ctl; NCKO spread across the circle but left gaps while Ctl cells formed a cohesive monolayer. Orange arrows indicate differential border patterning; purple arrows indicate gaps. (F) Live imaging for 48 h shows reduced spreading and increased post-division clustering in NUKO cells relative to Ctl and NCKO cells. Data are shown as mean +/− SEM (n=3 biological replicates; Student’s t-test). (G) IF of 8-day colonies stained for H3K27me3 (border associated), Sox2 (non-border), and phalloidin (F-actin). All genotypes establish concentric organization, indicating that *Nanog* 3’UTR deletion impairs spreading behavior without abolishing spatial organization. Scale bar (yellow, bottom right); A;150 um, C; 8.3 mm, D; 110 um, inset 25 um, E; left, 250 um, middle 125 um, right, 65 um, F; 15 um, G; left, 120 um, right, 40 um.

NCKO cells lacked detectable Nanog protein by western blot and immunofluorescence while retaining *Nanog* 3’UTR expression (Figure 3A; middle and right; Figure 3-figure supplement 1A,B). In contrast, NUKO cells retained intact *Nanog* CDS and normal Nanog protein expression but lacked the *Nanog* 3’UTR. (Figure 3A; middle and right, Figure 3-figure supplement 1A). These lines therefore uncouple Nanog protein expression from *Nanog* 3’UTR expression.

All lines express both Sox2 and Oct4 protein and remain alkaline phosphatase positive when maintained under pluripotent culture conditions (Figure 3B,C), indicating preservation of pluripotent identity. Under standard culture conditions, NUKO colonies were significantly smaller and more circular than either Ctl or NCKO colonies (Figure 3D).

When cultured for 72 hours on 500 um circular micropatterns, Ctl cells spread uniformly across the patterned area, whereas NCKO cells spread broadly but frequently left gaps in the monolayer (Figure 3E; purple arrows). In contrast, NUKO cells formed compact colonies that failed to spread cohesively across the patterned surface, resulting in irregular colony boundaries and lacking the smooth border arcs observed in Ctl and NCKO cultures (Figure 3E; orange arrows).

Live-cell imaging revealed marked differences in cell behavior following division. Ctl and NCKO cells extended protrusions and adhered as individual cells, whereas NUKO cells remained rounded, lacked extended protrusions, and frequently accumulated atop one another, resulting in significantly reduced surface spreading (Figure 3F). Thus, while both *Nanog* CDS and *Nanog* 3’UTR deleted cells remain pluripotent, loss of the *Nanog* 3’UTR is associated with pronounced defects in cell spreading and collective organization that are not observed following *Nanog* CDS deletion.

Because pluripotent ESCs normally exhibit concentric spatial patterning of *Nanog* mRNA domains and associated markers (Figures 1 and 2), we next asked whether this organization is retained following *Nanog* mRNA domain deletion. Ctl, NUKO, and NCKO cells were cultured as single colonies and stained for the border associated marker H3K27me3, the non-border marker Sox2, and F-actin (phalloidin). All three genotypes formed colonies with distinct H3K27me3 borders, an interior band of Sox2 expression and strong phalloidin staining in center cells (Figure 3G).

Notably, NUKO colonies consistently formed smaller more compact colonies with sharply defined borders, whereas Ctl and NCKO colonies spread more extensively, resulting in broader and less uniform spatial patterns. NCKO colonies frequently displayed an extended peripheral region of H3K37me3 positive cells (Figure 3G; yellow arrows). Together, these observations indicate that although *Nanog* 3’UTR deletion impairs cell spreading, the capacity to establish concentric organization is preserved.

### Transcriptional analysis distinguishes the effects of *Nanog* 3’UTR and *Nanog* CDS deletion in pluripotent ESCs

To assess the differential effects of *Nanog* 3’UTR vs *Nanog* CDS deletion, control (Ctl), *Nanog* CDS knockout (NCKO), and *Nanog* 3’UTR knockout (NUKO) mESC lines were compared by bulk RNA sequencing. Principal component analysis revealed clustering primarily by genotype, with replicate lines grouping more closely with each other than with other genotypes (Figure 4A). Both NUKO and NCKO cells were transcriptionally distinct from Ctl cells and from each other.

**Figure 4.**
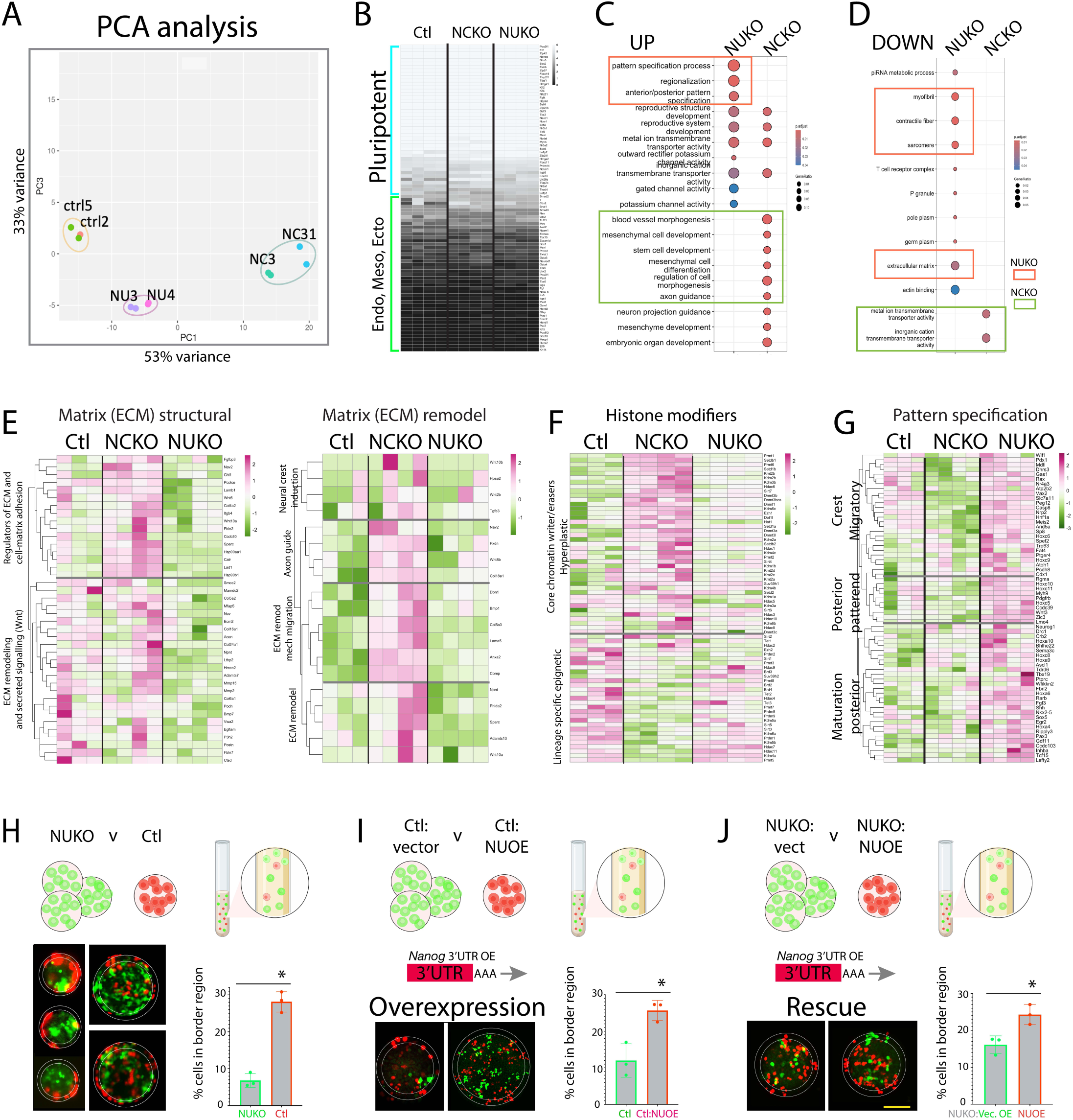
*Nanog* 3’UTR and *Nanog* CDS deletions uncouple transcriptional and spatial behaviors in pluripotent ESCs. (A) Principal component analysis (PCA) of bulk RNA-seq showing separation of control (Ctl), NUKO, and NCKO lines, with independent clonal replicates clustering by genotype. (B) Heatmap of pluripotency and differentiation associated gene expression showing broad retention of pluripotency markers across all lines. (C,D) Gene Ontology (GO) analysis of genes differentially expressed in NUKO and NCKO cells relative to Ctl, highlighting distinct biological processes associated with genes upregulated (C) or downregulated (D) in each genotype. NUKO cells show enrichment for processes related to early extracellular matrix (ECM) remodeling and pattern specification, whereas NCKO cells are enriched for developmentally associated and morphogenetic gene categories. (E) Heatmaps of ECM structural and ECM remodeling genes showing reduced expression of early ECM structural genes in NUKO cells and increased expression of ECM-related remodeling genes associated with epithelial organization and matrix stabilization in NCKO cells. (F) Heatmap of chromatin modifying gene expression showing broad alterations in NCKO cells, including reduced expression of TET family dioxygenases, compared with Ctl and NUKO cells. (G) Heatmap of pattern specification associated genes showing increased expression in NUKO cells relative to Ctl and NCKO cells, while pluripotency marker expression remains broadly maintained. (H) Mixed label assay comparing Ctl (red) and NUKO (green) cells showing preferential localization of Ctl cells to colony border regions. (I) Mixed label assay comparing Ctl cells and Ctl cells overexpressing the *Nanog* 3’UTR (Ctl:NUOE), showing enrichment of *Nanog* 3’UTR overexpressing cells at colony borders. (J) Mixed label assay comparing NUKO cells and NUKO cells overexpressing the *Nanog* 3’UTR (NUKO:NUOE), showing preferential localization of *Nanog* 3’UTR overexpressing cells to border regions. Data represent mean +/− SEM (n=3 biological replicates; Student’s t-test. Heatmap analysis for curated genes according to references in methods. Scale bar (yellow, bottom right); H; left,70 um, right, 100 um, I; left, 45 um, right, 120 um, J; left, 45 um, right, 45 um.

Neither NCKO nor NUKO lines showed broad downregulation of pluripotency associated genes or widespread upregulation of tissue-specific differentiation markers (Figure 4B). Consistent with this, all lines retained alkaline phosphatase activity (Figure 3C), and cells from each genotype were capable of integrating into host blastocysts (Figure 4-figure supplement 1A), indicating retention of pluripotent characteristics under the conditions examined.

Differential gene expression and Gene Ontology (GO) analyses revealed marked and distinct transcriptional differences between NUKO and NCKO cells relative to Ctl cells and to each other (Figure 4C,D). Prominent among these differences were changes in genes associated with cell adhesion and ECM organization. In NUKO cells, genes involved in early ECM remodeling processes, including matrix turnover and adhesion, were significantly downregulated (Figure 4E, left). In contrast, NCKO cells showed increased expression of distinct ECM-related and signaling genes associated with epithelial organization and matrix stabilization, including components of the WNT and Activin-associated pathways (Figure 4E, right).

In addition to ECM related changes, NCKO cells showed broad alterations in the expression of chromatin modifying enzymes, including reduced expression of TET family genes (Figure 4F), consistent with perturbation of chromatin associated regulatory programs following loss of Nanog protein. In contrast, NUKO cells displayed increased expression of genes associated with pattern specification, with enrichment for trunk, and posterior, neural associated gene programs (Figure 4G).

### *Nanog* 3’UTR influences border positioning independently of *Nanog* CDS expression

To assess whether *Nanog* 3’UTR expression affects colony cell positioning independently of the *Nanog* CDS, we performed gain-of-function mixing experiments in circular micropatterned cultures. Cells labeled with TdTomato or GFP were mixed and cultured for 72 hrs.

When Ctl cells (red) were mixed with NUKO cells (green), Ctl cells preferentially occupied colony border positions (Figure 4H). In contrast, when Ctl cells were mixed with Ctl cells overexpressing the *Nanog* 3’UTR (Ctl:NUOE), *Nanog* 3’UTR overexpressing cells were significantly enriched at colony borders relative to Ctl cells (Figure 4I). Similarly, when NUKO cells were mixed with NUKO cells overexpressing the *Nanog* 3’UTR (NUKO: NUOE), *Nanog* 3’UTR overexpressing cells again preferentially localized to border regions (Figure 4J).

Together, these findings indicate that increased *Nanog* 3’UTR expression-whether endogenous or ectopic-is associated with enhanced competitive occupancy of colony border regions in circular culture.

### *Nanog* 3’UTR loss alters morphogenetic behavior through ROCK-mediated cytoskeletal pathways

To determine whether the NUKO phenotype persists in the absence of the *Nanog* CDS, we generated full length *Nanog* mRNA KO lines (FLKO) (Figure 5; Figure 5-figure supplement 1A,B). Bulk RNA-seq analysis, migration associated marker expression and functional assays revealed that FLKO cells largely recapitulated the NUKO phenotype, while retaining some features associated with NCKO cells. Consistent with this, top downregulated categories include ameboidal migration and cell substrate adhesion (Figure 5-supplemental figure 5C). Further of the 40 extracellular matrix and remodeling genes downregulated in NUKO cells, more than half were also downregulated in FLKO cells (Figure 5-figure supplement 1D). In contrast, many ECM genes upregulated following *Nanog* CDS deletion were not similarly upregulated in FLKO cells (Figure 5-figure supplement 1E).

**Figure 5.**
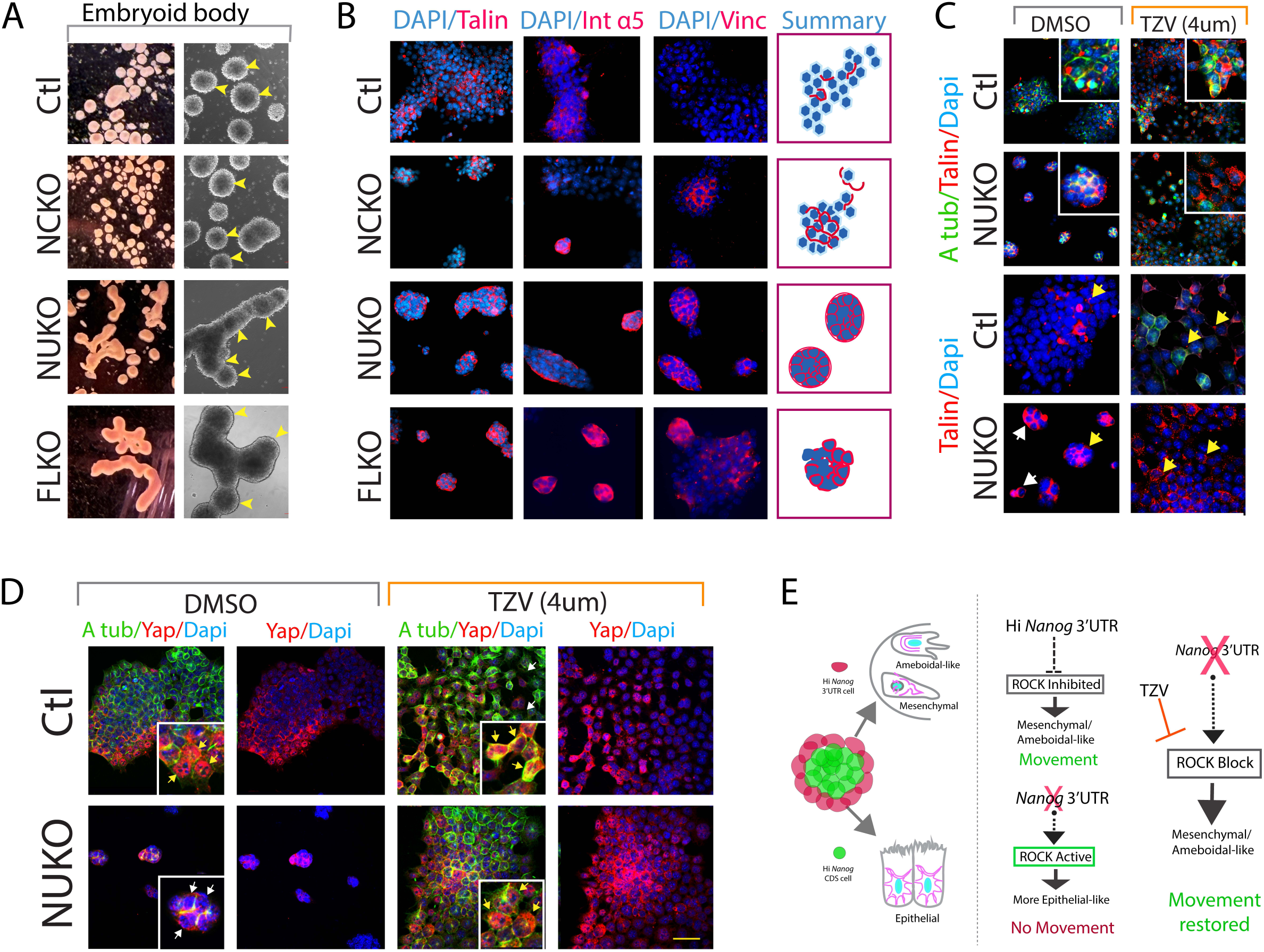
ROCK inhibition modulates cytoskeletal organization in *Nanog* 3’UTR deleted cells. (A) Embryoid body (EB) formation from Ctl, NCKO, NUKO and full-length knockout (FLKO) mESCs cultured for 2 days under non-adherent conditions. Ctl and NCKO cells form predominantly spherical EBs, whereas NUKO and FLKO cells form elongated, chainlike structures. Yellow arrowheads show “spheres”. Validation of FLKO lines in Figure 5-figure supplement 1A-E. (B) IF staining for cytoskeletal and adhesion-associated proteins, including Talin, integrin a5, and Vinculin, showing altered localization patterns in NUKO and FLKO compared to Ctl and NCKO cells. Summary schematics indicate representative protein distributions. Additional markers in Figure 5-figure supplement 1A-E; Supplementary Table 2. (C) Colony morphology and Talin localization in Ctl and NUKO cells treated with vehicle (DMSO) or the ROCK inhibitor Thiazovivin (TZV, 4μM). TZV treatment alters colony morphology and Talin distribution in NUKO cells. Insets show representative colonies. (D) IF staining for Yap in Ctl and NUKO cells treated with DMSO or TZV. YAP localization is predominantly cytoplasmic (white arrows) in NUKO and shifts following (yellow arrows) TZV (4 um) treatment. NCKO and FLKO cells shown in Figure 5-figure supplement 1H. (E) Schematic summary illustrating colony level morphological and cytoskeletal changes associated with *Nanog* 3’UTR deletion and their modulation by ROCK inhibition. Scale bar (yellow, bottom right); A; left, 980 um, right, 300 um, B; 60 um, C; 90 um, inset 30 um, D; 55 um, inset 25 um.

Morphogenic behavior was next assessed using embryoid body (EB) formation. After two days in non-adherent culture, Ctl and NCKO cells formed predominantly spherical EBs, (Figure 5A). In contrast both NUKO and FLKO cells formed elongated, tube-like structures composed of fused-spheroid chains, consistent with impaired remodeling or separation during EB formation (Figure 5A).

To further examine adhesion and cytoskeletal regulation, we assessed the expression and localization of adhesion and cytoskeleton-associated proteins including integrins, actin regulators and mechanical response proteins (Supplemental Table 2). Compared with Ctl cells, both NUKO and FLKO cells displayed altered localization of these proteins, characterized by uniform enrichment at colony borders (Figure 5B; Figure 5-figure supplement 1F). These patterns were largely absent in Ctl and NCKO cells.

Because cell migration and cytoskeletal organization are regulated by Rho GTPases-ROCK signaling, we tested whether pharmacological ROCK inhibition modifies the NUKO and FLKO phenotypes. Treatment with the ROCK inhibitor thiazovivin (TZV) altered colony morphology and normalized adhesion and cytoskeletal protein localization in both NUKO and FLKO cells (Figure 5C-E; Figure 5-figure supplement 1G-I). Consistent with this response, Talin-normally low in Ctl and NCKO cells-was elevated at cell junctions and colony edges in NUO and FLKO cells and was reduced following TZV treatment (Figure 5C, Figure 5-figure supplement 1G,I). Similarly, YAP localization, which was predominantly cytoplasmic in NUKO and FLKO cells, shifted to the nucleus following TZV treatment, a pattern normally observed in Ctl cells (Figure 5D; Figure 5-figure supplement 1H). Together, these findings indicated that *Nanog* 3’UTR loss is associated with altered ROCK-dependent cytoskeletal and adhesion behaviors.

### *Nanog* CDS and 3’UTR deletions differentially influence differentiation associated outputs

We next examined the contribution of *Nanog* mRNA components to maintenance of pluripotency and differentiation competence. Cells were subjected to eight days of leukemia inhibitory factor (LIF) withdrawal and assessed for colony integrity. Under these conditions, FLKO cells differed from NCKO, NUKO or Ctl cells, with the latter three maintaining more robust colony size and organization (Figure 6A). In contrast, FLKO colonies were largely depleted, suggesting that loss of both *Nanog* domains is more detrimental to colony maintenance that deletion of either the CDS or 3’UTR alone. Extended passaging under LIF-depleted conditions resulted in compromised colony structure across all genotypes, with FLKO colonies exhibiting the poorest survival.

**Figure 6.**
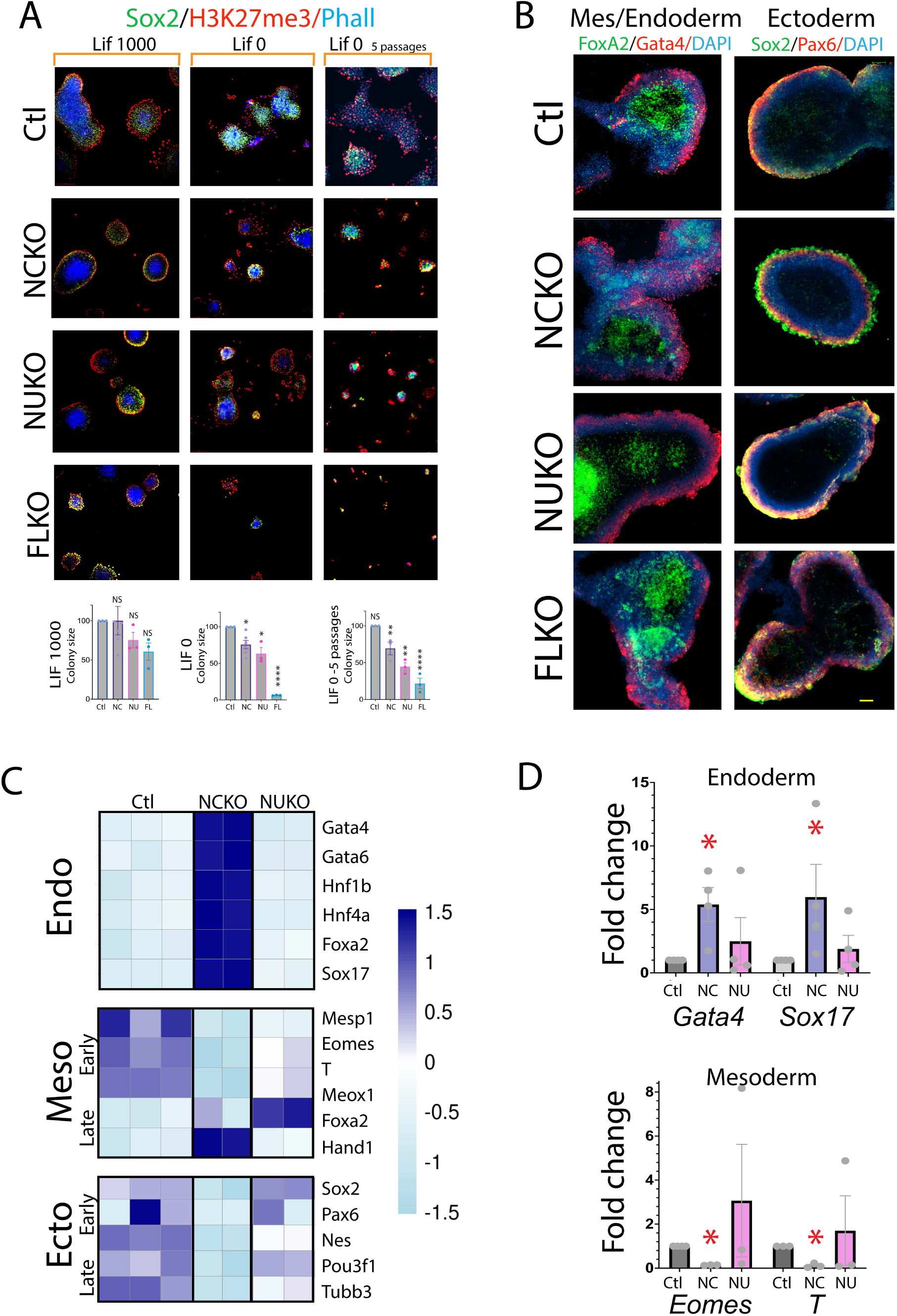
*Nanog* CDS and *Nanog* 3’UTR deletions differentially bias differentiation trajectories. (A) LIF withdrawal assays in Ctl, NCKO, NUKO and FLKO mESCs cultured without LIF for 8 days (left and middle columns) or following extended passaging under LIF depleted conditions (right column). Representative images are shown. Colony size was quantified. Data represent +/− SEM (n=3 biological replicates; > 5 colonies/replicate per condition; Student’s t-test). (B) Embryoid body differentiation followed by explant plating (10 days) showing expression of mesoderm/endoderm markers (FoxA2, Gata4) and ectoderm markers (Sox2, Pax6), indicating retained competence to generate derivatives of all three germ layers across genotypes. (C) Heatmap of bulk RNA-seq from plated 10-day differentiation cultures showing expression of representative endodermal (Endo), mesodermal (Meso) and ectodermal (Ecto), markers in Ctl, NCKO and NUKO cells. (D) qPCR analysis of sister cultures showing significant changes in representative endodermal (Gata4, Sox17) and mesodermal (Eomes, T) markers in NCKO cells relative to Ctl and NCKO cells. Additional markers tested did not show significant differences (Supplemental Table 3). Scale bar (yellow, bottom right); A; 160 um, B; 100 um.

Differentiation assays showed that all lines retained the capacity to generate cells expressing markers from each of the three germ layers, indicating preserved lineage competence (Figure 6B). Bulk RNA-seq analysis of plated 10-day differentiation cultures showed that NCKO cells exhibited pronounced positive endodermal bias (Figure 6C) with increased expression of endoderm associated genes and reduced expression of some early mesodermal markers (Figure 6C). These transctiptional shifts in endoderm were not observed in NUKO cells.

Quantitative PCR analysis of sister differentiation cultures confirmed significant changes in representative endodermal and mesodermal markers in NCKO cells, whereas corresponding changes in NUKO cells were not significant (Figure 6D; Supplementary Table 3). FLKO cells were capable of generating derivatives of all three germ layers, however consistent lineage bias could not be assessed due to limited survival of these cells under dissociated differentiation conditions.

Together, these findings indicate that deletion of either the *Nanog* CDS or 3’UTR preserves germ layer competence, but that these mRNA components differentially influence exit from pluripotency.

## Discussion

Nanog is essential for early embryogenesis yet dispensable for embryonic stem cell (ESC) self-renewal under standard culture conditions, leading to a longstanding paradox about its developmental function. Here, we provide a framework for understanding this paradox by showing that distinct regions of the *Nanog* RNA encode separable regulatory activities. Differential usage of the *Nanog* coding sequence (CDS) and 3’untranslated separates gene output into distinct functional contributions, revealing an RNA-based regulatory layer that extends Nanog function beyond protein output alone and helps explain why Nanog is required during development but not under standard ESC culture conditions.

Deletion of the *Nanog* 3′UTR produced marked defects in cell spreading, extracellular matrix remodeling, and cytoskeletal organization, accompanied by increased cellular aggregation and failure to form cohesive monolayers. These phenotypes correlated with altered localization of focal adhesion and actin-associated proteins and were rescued by pharmacological inhibition of ROCK kinase signaling, placing the *Nanog* 3′UTR functionally upstream of canonical migration and adhesion pathways. Across experimental contexts, including standard culture, micropatterned colonies and embryoid body formation, cells enriched for *Nanog* 3’UTR expression preferentially occupied colony borders, whereas cells lacking the 3′UTR formed compact colonies and failed to form discrete spheroids. Together, these observations identify an unrecognized role for the *Nanog* 3’UTR in regulating cell movement and early morphogenetic behaviors.

In contrast, deletion of the *Nanog* CDS produced a distinct transcriptional phenotype characterized by upregulation of ECM and WNT associated gene programs linked to epithelial polarization and colony sculpting, along with widespread changes in chromatin modifying enzymes. Reduced expression of TET family dioxygenases and altered balance of chromatin writers and erasers are consistent with perturbation of chromatin regulatory programs. These findings support a canonical role for the Nanog protein in maintaining epithelial restraint and appropriate chromatin plasticity, thereby preserving a balanced pluripotent state. Thus, while the *Nanog* CDS primally contributes to transcriptional and epigenetic regulation, the *Nanog* 3’UTR exerts an independent influence on cell behavior and tissue organization. Our results reinforce the established role of Nanog protein in pluripotency, while revealing an additional RNA-based regulatory function for the *Nanog* transcript in cell morphogenesis.

Analysis of full-length *Nanog* knockout (FLKO) cells further supports an instructive role for the *Nanog* 3’UTR that cannot be explained by loss of the CDS alone. Across adhesion and ECM associated gene programs, FLKO transcriptional profiles more closely resemble those of *Nanog* 3’UTR knockout cells than those of *Nanog* CDS knockout cells. Similarly, FLKO cells share cytoskeletal colony organization defects with *Nanog* 3’UTR deleted cells and fail to form discrete spheroids in three-dimensional cultures. These findings argue that the morphogenetic phenotypes associated with *Nanog* 3’UTR loss reflect a direct contribution of this RNA region to cell behavior rather than secondary effects of altered protein activity.

Differential loss of *Nanog* mRNA domains also influences differentiation trajectories. While neither the CDS nor the 3’UTR is strictly required for germ layer competence, the two domains shape exit trajectories in distinct ways. Deletion of the *Nanog* CDS biases differentiation toward endoderm associated programs, whereas deletion of the *Nanog* 3’UTR does not. The increases in pattern specification genes observed in *Nanog* 3’UTR deleted cells are not associated with detectable changes in lineage. Together, these observations further support the idea that *Nanog* mRNA domains encode separable regulatory outputs that influence developmental progression.

Micropatterned culture experiments reveal that ESCs self-organize into concentric regions defined by *Nanog* mRNA domain usage, with *Nanog* 3’UTR enriched cells preferentially located at colony borders and *Nanog* CDS enriched cells residing more internally. Although both populations remain pluripotent, this spatial arrangement suggests functional specialization within the colony, with border cells preferentially executing morphogenetic behaviors and interior cells contributing to epithelial organization and mature colony sculpting. These findings indicate that pluripotent colonies can exhibit coordinated functional specialization, mediated, at least in part, by differential mRNA component usage.

The mechanisms that generate isolated 3’UTR or CDS mRNA components remain incompletely understood and may involve differential stabilization, alternative polyadenylation or post-transcriptional processing. While our study does not resolve these mechanisms, it establishes functional significance for mRNA components themselves and highlights RNA domain usage as a biologically meaningful layer of regulation. Isolated 3′UTRs share structural features with long noncoding RNAs, yet their origin in protein-coding transcripts provides additional contextual information that may aid in interpreting their potential roles. The accompanying lookup table provides a resource for examining 3’UTR/CDS usage across additional genes, enabling broader exploration of mRNA domain-specific regulation.

In summary, we identify distinct and complementary functions for the *Nanog* 3’UTR and *Nanog* CDS, supporting a previously underappreciated mode of gene regulation operating through mRNA architecture. By demonstrating that different regions of a single transcript can encode separable cellular behaviors, this work expands the concept of gene function beyond protein output and underscores the importance of RNA-based regulatory mechanisms in pluripotent cells and early development.

## Materials and Methods

### Mouse and human embryos

Human blastocysts originated from the Stanford Fertility and Reproductive Health Center in excess of reproductive need. The blastocysts were donated to the RENEW Biobank at Stanford University (IRB10466) after In Vitro Fertilization, embryo culture and vitrification on Day 5 or 6. As approved by a SCRO protocol (SCRO-795), a total of 16 human vitrified blastocysts were warmed using Kitazato vitrification/warming protocol, allowing 1-2hrs for re-expansion prior to them being fixed. Mouse embryos were obtained from Stanford transgenic, knockout and tumor model center. Briefly, 50-80 mouse zygotes were made each time from in vitro fertilization (IVF) and transferred to 30 µl drops of culture medium covered with sterile mineral oil on a culture dish and placed in culture incubator at 37°C under 5% CO2 until reaching the morula or blastocyst stages. ISH and antibody stain was carried out with methods described above with the modification that each embryo was processed within a series of single droplets of 15ul hybridization and wash solutions while situated on a microscope slide.

For mouse blastocyst injection, Ctl, NCKO and NUKO mESCs were labelled with lentivirus expressing tdTomato, and then injected into the mouse blastocyst, cultured in embryo culture medium for 48 hours before paraformaldehyde fixation. The percent of cells for each line with residency in the ICM was measured by visual inspection. For blastocyst integration assays, two independent NCKO and NUKO mESC clones were used. Cells were labeled with tdTomato via lentiviral transduction, injected into mouse host blastocysts, cultured for 48h with KSOM/ Embryo-Love medium in culture incubator at 37°C under 5% CO2, fixed, and assessed for localization with the ICM by microscopy. ICM residency was scored as the number of fluorescent donor cells localized to the ICM per embryo.

### mESC and hESC Cell culture

Strain 129 mouse embryonic stem cells (mESCs) were obtained from the lab of Dr. Tessier-Lavigne (https://www.ncbi.nlm.nih.gov/pmc/articles/PMC2747754/), cultured in E14 plus complete medium (Glasgow Minimum Essential medium (Sigma-Aldrich G5154-6) supplemented with 10% heat inactivated FBS (Fisher Scientific SH3007103), 0.1 mM non-essential amino acids (Thermo Fisher Scientific 11140050), 1 mM sodium pyruvate (Thermo Fisher Scientific 11360070), 1x Pen/Strep (Thermo Fisher Scientific 15140122), 1x β-mercaptoethanol (55 um) (Thermo Fisher Scientific 21985023), 1,000 units/ml Leukemia Inhibitory Factor (LIF) (Millipore Sigma ESG1106), and freshly added Glutamax (Thermo Fisher Scientific #35050061) (final concentration of 1X), 1 um PD03259010 (Reprocell 04-0006-10) and 3 um CHIR99021 (Cellagen Technology C2447-50)), and used within one week. mESCs were maintained and split a 1:6 ratio with 2x trypsin (0.8% Trypsin (Invitrogen) in PBS with EDTA-trisodium salt and chicken serum) for 2 minutes at 37°c every two days, and re-plated in 6 well plates that had been precoated with 0.1% gelatin (Sigma Millipore ES-006-B) for 30 minutes at 37’C. For embryoid body culture, cells were plated in low affinity 6-well plate with a density of 5×10 cells/ml in mESC EB diff medium (DMEM with10% regular FBS+ Non-essential amino acids+β-mercaptoethanol+Pen/Strep). H7 human embryonic stem cells (hESCs) were incubated at 37 C, 5% CO2 in mTeSR1Plus (STEMCELL Technologies 85850) complete medium (mTeSR1 basal medium with supplement made according to manufacturer’s protocol). hESCs were obtained from WiCell (RRID:CVCL_9772), maintained and split from around 90% confluence with 1x Versene (Fisher Scientific 15-040-066) for 5 minutes at RT and then split 1:8 into 6 well plates precoated with 1:200 diluted geltrex in DMEM/F12 medium (Thermo Scientific A1413201) for 30 minutes at 37’C. All cells were incubated at 37 C with 5% CO2.

### Fluorescence in situ hybridization (ISH)

Two color fluorescence in situ hybridization performed as previously described (Kocabas, Duarte et al. 2015, Ji, Yang et al. 2021)(Ji *et al*., 2021; Kocabas *et al*., 2015), using the TSA Plus Cyanine 3 & Fluorescein (NEL753001KT, PerkinElmer, Waltham, MA) kit according to the manufacturer’s instructions. Briefly, cells in culture were fixed with 4% paraformaldehyde (PFA) in PBS for 15 min, treated with proteinase K (1ug/ml) for 5 min (RT), acetylated (1% triethanolamine and 0.25% acetic anhydride) for 10 min, pre-hybridized (50% formamide, 5X SSC, 5X Denhardt’s solution.5 mg/ml Herring sperm DNA and 250 μg/ml Yeast tRNA) for 10 min, then hybridized with fluorescein labeled CDS and Digoxygenin (DIG) labeled 3’UTR probes, diluted in prehybridization buffer (0.5 ng/ul), and hybridized at 56°C for 3 hours. Proper washes were carried out accordingly between each step. Specimens were then sequentially post-stained with fluorescein or Cy3 chromogens respectively, and the corresponding Alexa Fluor 647 secondary antibodies and Hoechst 33342 for DNA staining and mounted with Fluoromount-G mounting medium (Southern Biotech catalog #0100-01). The ISH probes are listed in Supplemental Table 1.

### Micropattern cell culture

Micropattern CYTOO™ (Arena A,™, France) chip cell culture was performed as previously described by us and others (Warmflash, Sorre et al. 2014, Morgani, Metzger et al. 2018, Ji, Yang et al. 2021). Patterns of 1000, 500, 250, 120 and 80 um circles are dispersed over the full surface. Briefly, for mESC, 700 ul drops of 20 ug/ml mouse Laminin (Sigma, L20202) in PBS without calcium and magnesium (PBS−/−) was spotted onto parafilm sitting in a 15 cm tissue culture plate. After one wash in PBS−/−, chips were inverted on top of the drops for 2h at 37°C. They were then washed 5x with PBS−/− and a single cell suspension of mESC cells (2 × 10^6^) was evenly plated onto chips grown in 6-well plates for three days in complete E14 medium before fixation and ISH. For hESC, 700 ul drops of 5 ug/ml human laminin (LN521) (Biolamina) in PBS with calcium and magnesium (PBS+/+) in were spotted onto parafilm in a 15 cm tissue culture plate. After one wash in PBS+/+, chips were inverted on top of the drops for 2h at 37°C, then washed 5x with PBS+/+ and a single cell suspension of mESC cells (2 × 10^6^) was evenly plated onto chips, grown in 6-well plates for 72h in complete mTeSR medium before fixation and ISH. Quantitation for the number of high *Nanog* 3’UTR (red or high CDS cells (green) in aggregates, or alone (less than 3 cells), was measured across four independent field for three experiments. Statistical significance was determined by Chi-square analyses (X^2^). Similarly, for the number of high *Nanog* 3’UTR cells, in CYTOO circles of different sizes red cells were counted and presented as a fraction of total DAPI positive cells. Statistics are the mean of means for three independent experiments for each condition, analyzed by Student’s t-test.

### Cas9 and CRISPR gRNAs knockout stable mESC cell lines

CRISPR knockout studies were carried out in mESCs, where Nanog is represented by a single expressed genotype, enabling straightforward targeting, while the human genome contains multiple Nanog pseudogenes possibly complicating interpretation.

gRNAs targeting different gene CDS, 3’UTR regions or CDS+ 3’UTR regions were designed individually using IDT Custom Alt-R CRISPR-Cas9 guide RNA design tool, specific gRNAs were selected for both their on-target potential score and off-target risk score. gRNAs were synthesized by IDT, gRNAs then were cloned into modified pL-CRISPR.EFS.GFP construct (RRID:Addgene, 57818) with removal of Cas9, and GFP was replaced by tdTomato for 3’ targeting gRNAs. 3’UTRs and Cas9 resistant rescue *Nanog* 3’UTR were cloned into pLKO-H2BmRFP1 construct, a generous gift from Dr. Fuchs; Rockefeller University. The Cas9 expression (RRID:Addgene 52962) and rtTA constructs (RRID:Addgene_61472) were obtained from Addgene.

A Cas9 mESC stable cell line was made by lentiviral infection of mESCs with lentiCas9-Blast, screened for single colonies by blasticidin treatment (10 ug/ml) and evaluated by PCR. Lentivirus was made according to the lentivirus production section, and two different gRNAs with a total of 2x10e7 IU, along with 5ug/ml polybrene (Sigma), were used to infect the cells for knockout/knockdown. Knockout efficiency was verified by PCR of genomic DNA with primers targeting the deleted region. For rescue experiments, equal amounts of Cas9 resistant *Nanog* 3’UTR virus, with 5x10e6 reverse tetracycline-controlled-activator (rtTA) virus, was delivered simultaneous with knockout gRNA viruses. *Nanog* CDS, 3’UTR or full length CRISPR gRNA knockout mES cell lines were produced by transient transfection of mESCs with Cas9 and corresponding gRNAs, single cells were then FACS sorted into 96-well plates after GFP or tdTomato selection (expressed on the same gRNA vectors) Single positive clones were obtained by genomic DNA PCR and western blot analysis. Two independent NCKO, NUKO and FLKO lines were generated and validated by genomic PCR, RNAseq IGV trace and Western blot for Nanog protein.

### Colony formation and LIF deprivation assay

Ctl and knockout cell lines were each plated at 1000 cells /24-well plate well and grown for five days in decreasing concentrations of Leukemia Inhibitory Factor (LIF): 1000 U/mL (standard), 100 U/mL, 30 U/mL, and 0 U/mL before fixation, imaging, and quantification. Differentiation for the various cell lines; control (C2, C5), NCKO (NC3,NC31), NUKO (NU3, NU4), and full-length knockout (FL58, FL84) was assessed by measuring surviving cells in each line. Colony integrity and differentiation were evaluated using immunofluorescence markers: H3K27me3 (edge cell marker), Sox2 (pluripotency marker), and Phalloidin (mid-colony). For Figure 5F, colony sizes were measured across three fields for each line. For 5G, as there were few to no colonies in the FLKO lines, therefore total fluorescence across three fields was measured for each line. Statistics are the mean of means for three independent experiments for each condition, analyzed by Student’s t-test.

### Alkaline phosphatase stain

Alkaline phosphatase (AP) staining was according to vendor instructions, briefly, cells were fixed with fixation solution for 5 minutes at RT, and then stained with freshly prepared AP substrate solution in the dark, for 10 minutes, RT. The reaction was stopped by washing the cells twice with 1× PBS.

### Cell mixture rescue assay

Different groups of mESCs were transfected with pLV-H2BRFP construct that containing a Cas9 resistant NGG mutation *Nanog* 3’UTR sequence or modified pLV-H2BRFP construct that replaced RFP with GFP. Differentially transfected GFP or tdTomato or tdTomato plus NUOE cells were mixed 9:1 and plated on CYTOO chips for 72h. Cells were then fixed, stained for anti-GFP (green secondary) or anti-tdTomato (red secondary) and quantified. Briefly, in post image analyses, CYTOO chip circles were covered with a preformed concentric grid and the percent of total green cells in the outer circle, compared to the percent of total red cells in the outer circle, and the ratio of red; green calculated. Statistics are the mean of means for three independent experiments for each condition, analyzed by Student’s t-test.

For mixed colony assays, GFP or tdTomato or with tdTomato expressing mESCs were mixed at 9:1 rations and plated on CYTOO™ micropatterned circles for 72h. For rescue, dTomato + NUOE (*Nanog* 3’UTR, CAS9-resistant) cells were used. After fixation and IF, concentric ring grids were applied, and the percentage of each population located in the out ring was measured and expressed as a ratio. Three biological replicates, each with > 8 technical replicate fields were quantified and statistical significance determined by Student’s t-test.

### Live imaging of Cultured Cells

mESCs were grown in 24-well plates in E14 plus complete medium. Time-lapse imaging was carried out using a computer-assisted fluorescence microscope (Nikon) equipped with an objective lens, LED and a CCD camera. For DIC and fluorescent imaging, images were acquired every 20 minutes for 24 hours using Nikon NIS-elements AR software, exported as mp4 video files and analyzed in Fiji (Fiji, RRID:SCR_002285). Cell spread was measured by the tissue culture plate area occupied across three divisions, measured using ImageJ (Schindelin, Arganda-Carreras et al. 2012). Statistics were from independent videos for each condition, analyzed by Student’s t-test.

Thiazovivin (Selleck Chemicals S1459) and Y-27632 are both ROCK inhibitors and showed similar effects. Additional agents tested that did not significantly affect migration include CNO3 (Rho activator II), Defactinib (FAK inhibitor), Forskolin (increase cyclic AMP), Ml141 (Cdc42 inhibitor), and Jasplakinolide (actin stabilizer). Cycloheximide was used in Nanog protein half-life western blot assay at 50 ug/ml for indicated time.

The dose response for each drug was based on the IC50 listed in manufactures specifications. With cultures treated for 24 hours.

### qPCR

RNA was extracted using the Qiagen RNeasy plus mini kit (Qiagen 74134) and 2 µg of total RNA was reverse transcribed to cDNA using the SuperScript IV (Thermo Fisher Scientific, 18091050) both according to manufacturer instructions. qPCR assays were performed using the LightCycler 480 II (Roche) in 10µL reaction volume containing 6µL of SYBR green Master mix (Fisher Scientific A25918), 2µL of diluted cDNA (20 ng) and 1µL (300 nmol) each of a gene-specific forward and reverse primer. The following standard PCR reaction conditions were used for all transcripts: 95 °C 2 min; 40 cycles of 95 °C 10 s, 60 °C 1 min; 1 cycle of 95 °C 15 s, 60 °C 15 s, 95 °C 15 s. Each PCR sample was carried out in quadruplicate and repeated three times to ensure reproducibility. For data analysis, we adopted the 2−delta threshold cycle (Ct) method. The Ct of each target gene was first normalized to the Ct of GAPDH in each sample and then to the corresponding value in the control sample and expressed as log2. Data shown is change normalized to Ctl line. 4 technical replicates, three biological replicates.

### Immunofluorescence Staining

Cells were fixed in 4% paraformaldehyde, 15 minutes, RT, washed 3x in PBS, permeabilized with 0.2% Triton X-100 in PBS for 10 minutes, and blocked using 3% BSA in PBST (PBS + 0.1% Tween-20) for 1 hour, RT. Primary antibodies were added at 1:500 or as per manufacturer’s recommendation in 1% BSA in PBST and incubated at 4’C overnight. Cells were then incubated with the corresponding Alexa Fluor secondary antibodies (1:1000 dilution) for 1 hour at room temperature in the dark. Nuclei were counterstained with (DAPI) (Thermo Fisher Scientific) at 1:5000 for 10 minutes. Coverslips were mounted onto Superfrost Plus glass slides (Invitrogen) using Fluoromount-G mounting medium (Southern Biotech).

### Lentivirus

Lentiviruses of the sgRNAs, rtTA, and pLV overexpression constructs were obtained by co-transfection of library plasmids with two viral packaging plasmids psPAX2 (RRID: Addgene_12260) and pMD2.G (RRID:Addgene_12259) into HEK293T using jetPRIME transfection reagents (VWR 89129-926) according to manufacturer’s protocol. Human embryonic kidney (HEK293) cell line, originally obtained from ATCC (ATCC Cat# CRL-3216, RRID:CVCL_0063), was grown in Dulbecco’s modified Eagle’s medium (DMEM;Thermo Fisher Scientific 11960069) supplemented with 10% fetal bovine serum (FBS; Gibco10437028), and 1X Penicillin Streptomycin Glutamine (Fisher Scientific 10-378-016)

### Bioinformatic analyses 3’UTR/CDS ratio quantification

Raw data for hESC and mESC scRNA seq are downloaded from GEO database under accession number GSE75748 (https://www.ncbi.nlm.nih.gov/geo/query/acc.cgi?acc=GSE75748) and ArrayExpress database (http://www.ebi.ac.uk/arrayexpress) under accession number E-MTAB-2600. Quantification of 3’UTR and CDS expression as described (Kocabas, Duarte et al. 2015, Ji, Yang et al. 2021). To obtain expression matrices for 3’UTR and CDS respectively, we quantified gene average coverage for 3’UTR or CDS separately using the bedtools (v2.17.0) function coveragebed with the alignment file (BAM). Genes with coverage ≤ 5 mapped reads of both CDS and UTR were removed. CDS or 3’UTR expression was normalized to their library size (total mapped reads) and quantified as transcripts per base pair. The ratio of 3’UTR/CDS is defined as normalized 3’UTR counts/ (normalized 3’UTR counts + normalized CDS counts) and is distributed from 0-1: 0 =CDS only; 1=3’UTR only. Genes with <5 mapped reads in both the 3’UTR and CDS regions were excluded.

### Gene ontology and functional categorization

Gene categories were initially assigned by GO analysis (Gene Ontology, Aleksander et al. 2023), defined as p-value ≤ 0.05 and absolute log2 fold change ≥ 0.05 Deseq2(Love, Huber et al. 2014), using clusterProfiler (v4.4.4) (Ashburner, Ball et al. 2000), focusing on Biological Process ontology. Results were then visualized using enrichplot.

This was followed by targeted manual curation to of specific gene expression within a GO category. Definitions for various cell types were informed by established literature describing; pluripotency (Shahbazi and Zernicka-Goetz 2018), ECM remodeling (Page-McCaw, Ewald et al. 2007, Legate, Wickstrom et al. 2009, Hynes and Naba 2012), ECM stabilization and epithelial organization (Li, Edgar et al. 2003, Arnold and Robertson 2009, Eiraku, Takata et al. 2011), pattern specification (Neijts, Simmini et al. 2014, Deschamps and Duboule 2017), and chromatin balance linked to epithelial organization (Orkin and Hochedlinger 2011, Kraushaar and Zhao 2013, Che, Lee et al. 2022), downregulation of TET dioxygenase activity linked to restriction of differentiation capacity (Dawlaty, Breiling et al. 2014, Zhang, Zhang et al. 2023).

### Read processing and mapping

RNAseq data were processed using the RNA-seq workflow in snakePipes (v2.5.3)(Bhardwaj, Heyne et al. 2019). Within the pipeline, reads were mapped against human (hg38) or mouse (mm10) reference genomes using STAR (v2.7.3a) with default parameters. Gene expression quantification was quantified using RSEM (v1.3.1) yielding in expected raw counts and TPM with default parameters.

### PCA Component Analysis (PCA)

PCA was performed on variance-stabilized gene expression RNAseq data using plotPCA function from the DESeq2 (v1.36.0). This step was used to assess the transcriptomic variance and visualize clustering across the samples. Variance stabilizing transformation (VST) was used prior to PCA to minimize the effect of sequencing depth and further improve interoperability of the principal components.

### Cell clustering

The R package “Seurat” was applied to analysis single-cell data(Satija, Farrell et al. 2015), excluding poor quality cells as described in (Kolodziejczyk, Kim et al. 2015, Chu, Leng et al. 2016). Cells are clustered by t-SNE with the number of clusters chosen automatically by Seurat.

### Website development

For an interactive visualization of gene-specific expression patterns and 3’UTR/CDS ratio data (3’UTR/(3’UTR plus CDS) reads) we developed a web-based application through ShinyApps framework. The application displays t-distributed stochastic neighbor embedding (t-SNE) plots from hESC and mESC scRNA-seq data which allows users to explore expression for any gene of interest. The application was deployed using R and <u>Shinyapps.io</u> platform which makes it publicly available for interactive exploration.

During the preparation of this work the authors used Chat GTP4 (OpenAI. ChatGPT. https://chat.openai.com (2024)) to increase brevity, clarity and conciseness and to gain additional insight into lists of RNA expression level changes After using, the authors reviewed and edited the content as needed and take full responsibility for the content of the published article.

## Supporting information

Yang supp 1-1

Yang supp 1-2

Yang supp 2-1

Yang supp 3-1

Yang supp 4-1

Yang supp 5-1

## Acknowledgements

We thank Subramaniyam Ravichandran, Mujib Fnu, Chenjie Pan for technical assistance and Marc Tessier-Lavigne, Richard Axel, Jeffrey Friedman, Sally Temple and Mary Beth Hatten and for critical reading of early versions of the manuscript.

## Author contributions

Z.Y.; conception and design, collection and/or assembly of data, data analysis and interpretation, manuscript writing, final approval of manuscript. S.J.; conception and design, collection and/or assembly of data, data analysis and interpretation, manuscript writing. P.K., K.I. M.S., L.G., and S.P.; collection and/or assembly of data, data analysis and interpretation. B.B.; provision of study material or patients, training.

Further information and requests for resources and reagents should be directed to and will be fulfilled by the lead contact, Mary Hynes.

This study did not generate new unique reagents.

