## Supplementary figures and images for "Functional Separation of mRNA Domains Coordinates Pluripotent Cell Behavior"

### Yang supp 1-1

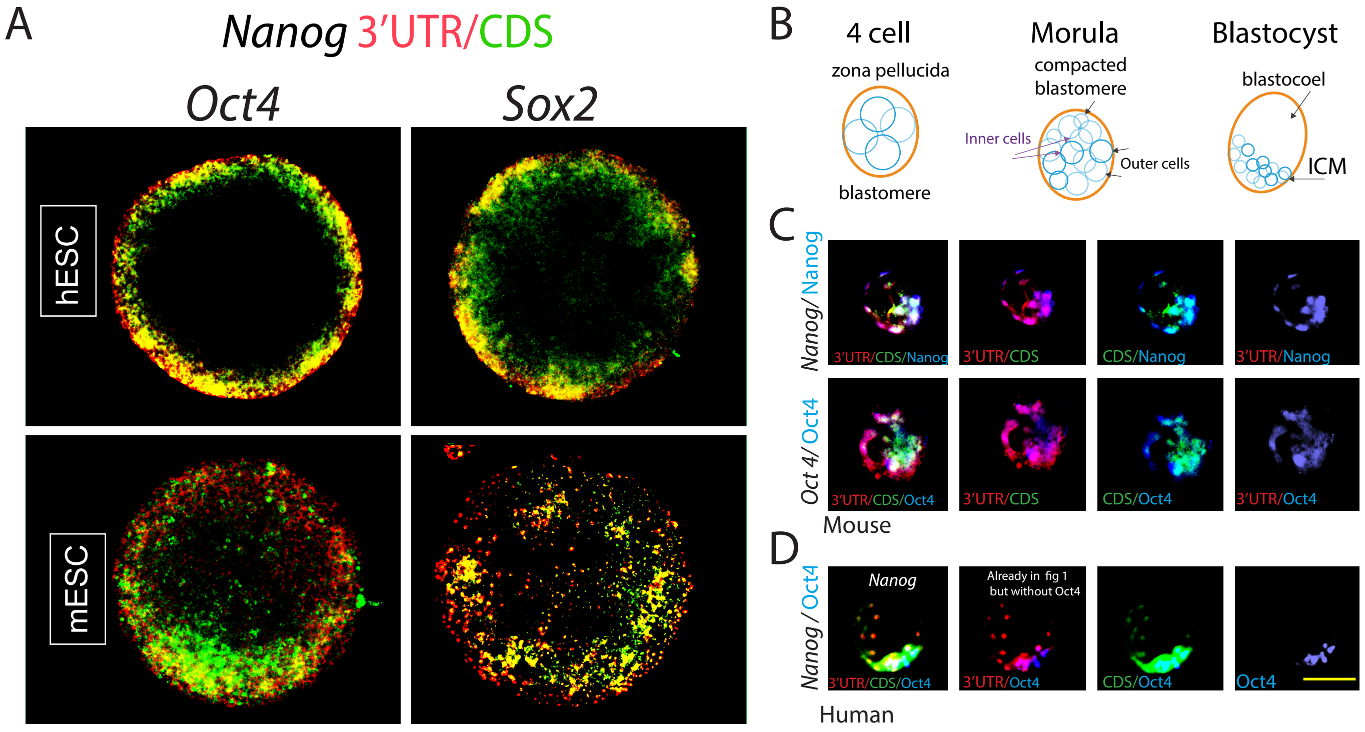

### Yang supp 1-2

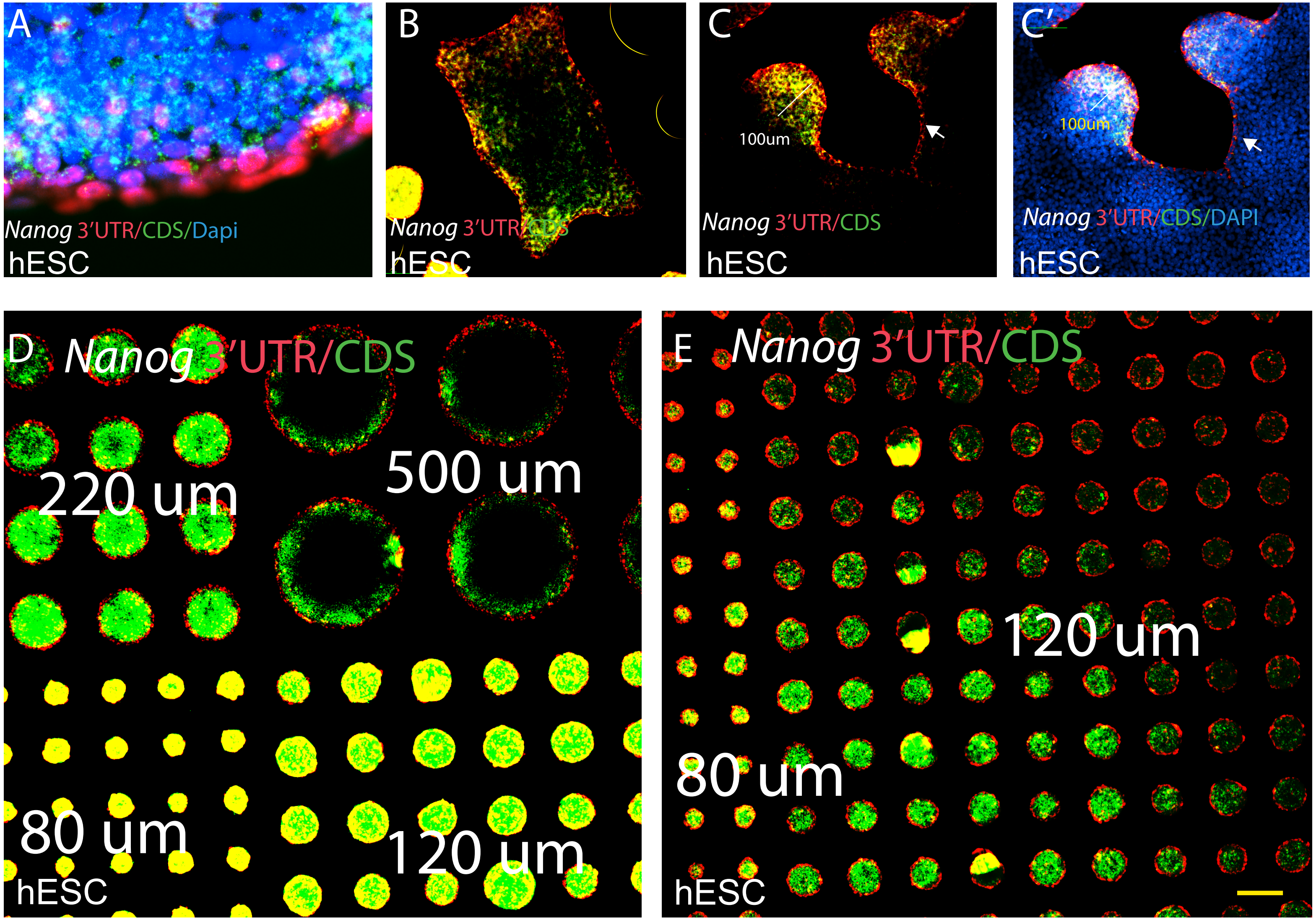

### Yang supp 2-1

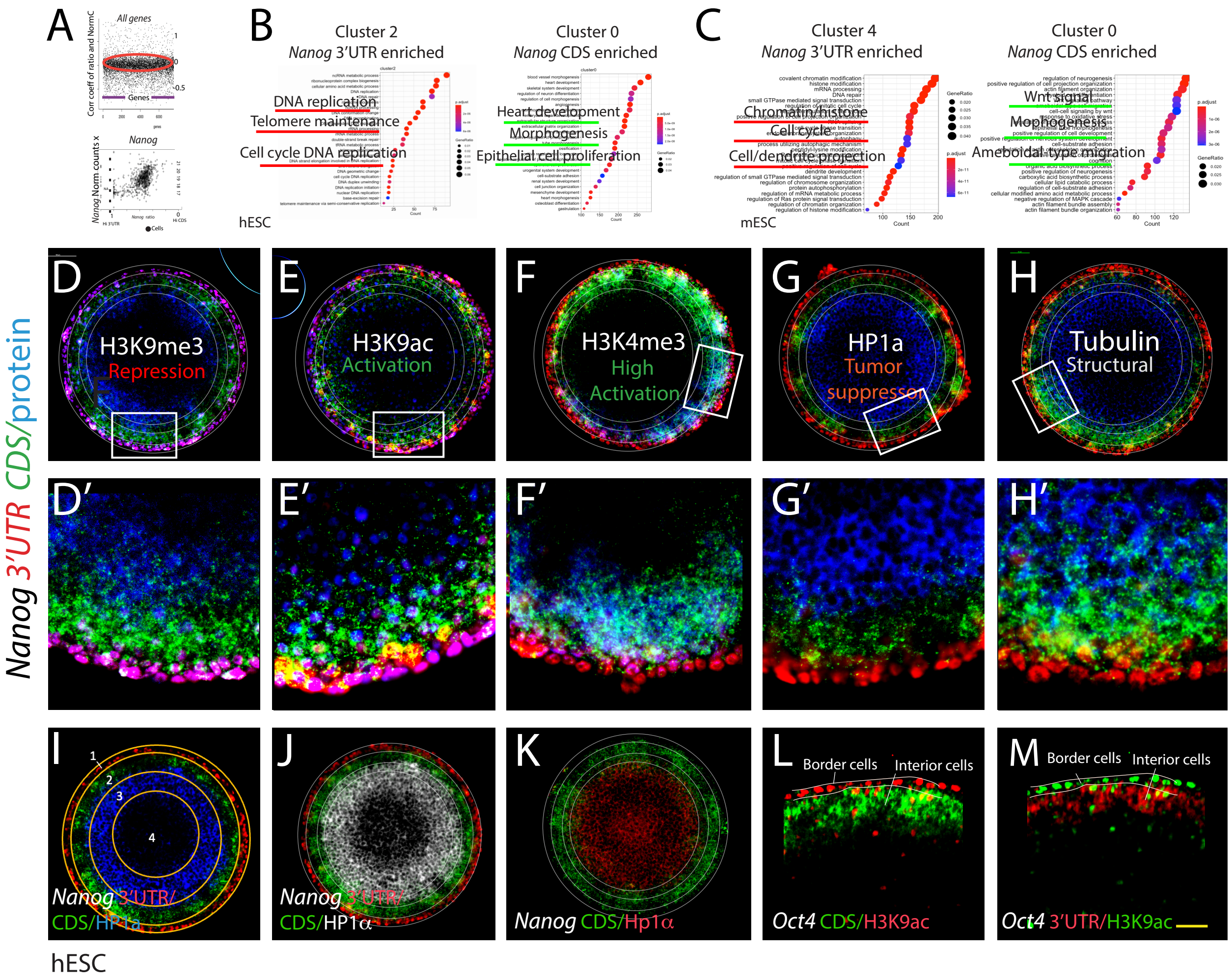

### Yang supp 3-1

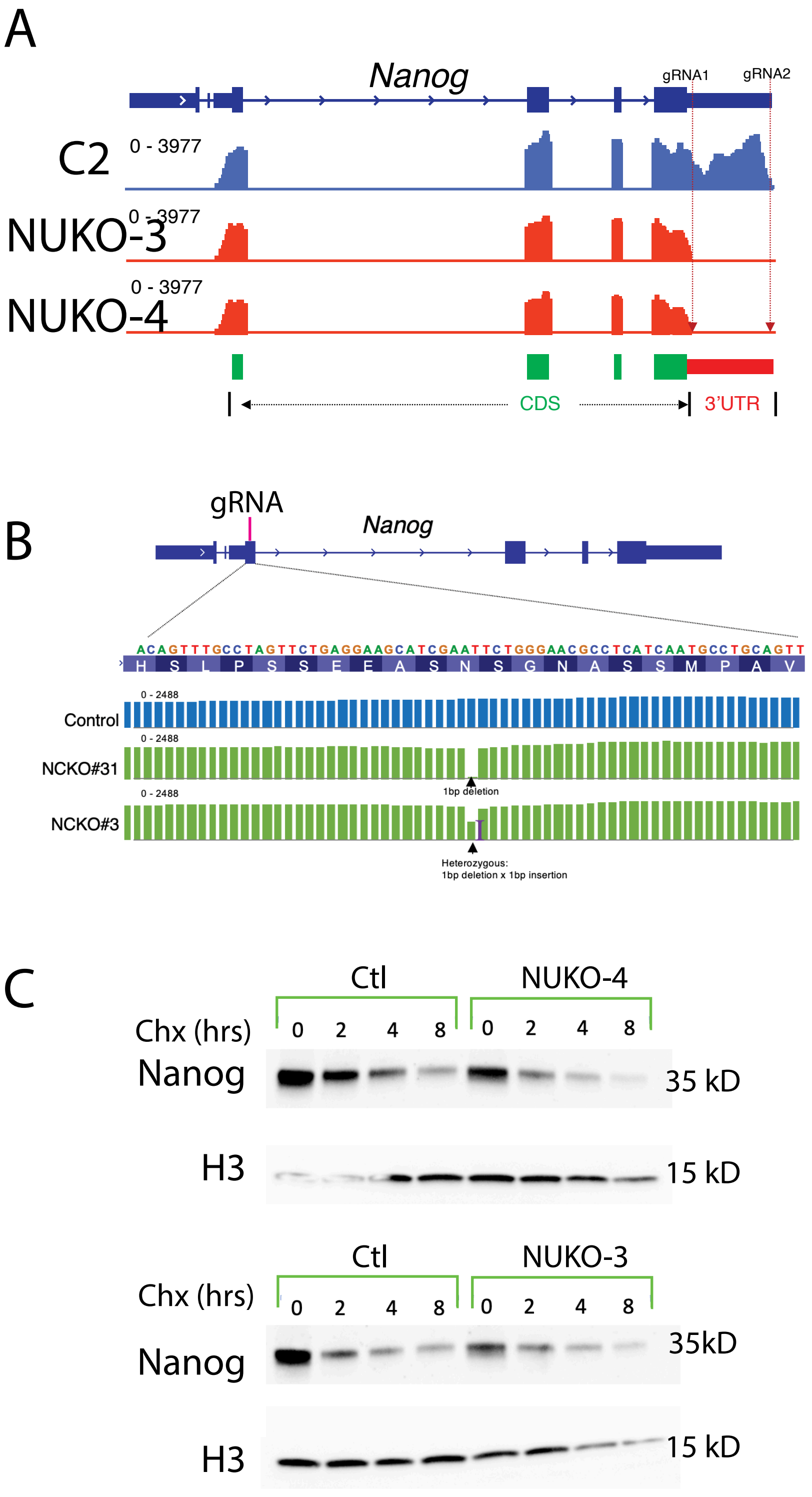

### Yang supp 4-1

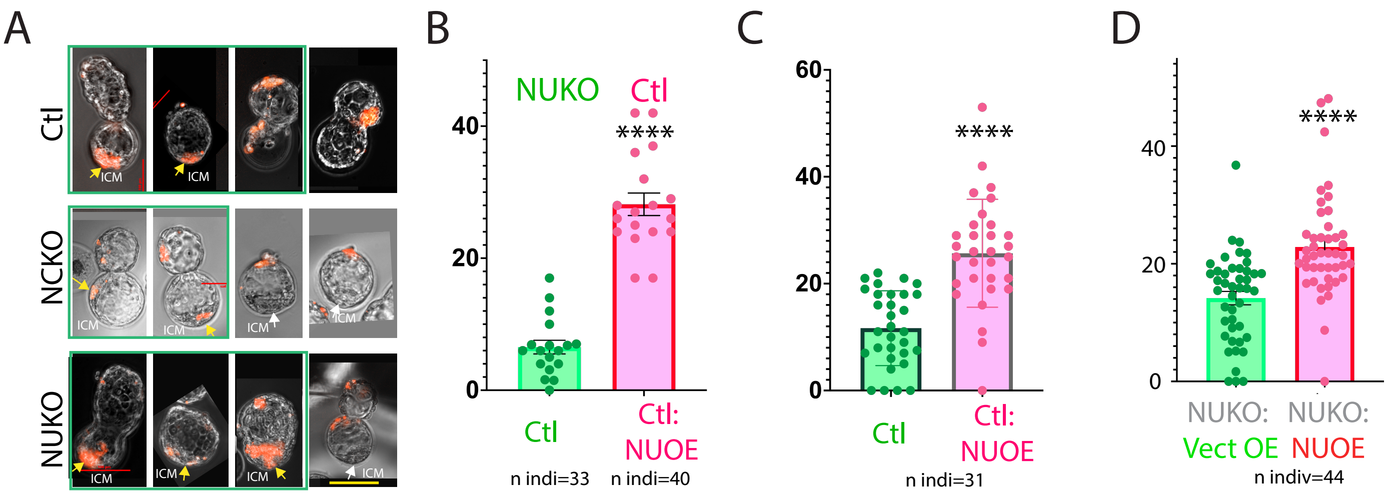

### Yang supp 5-1

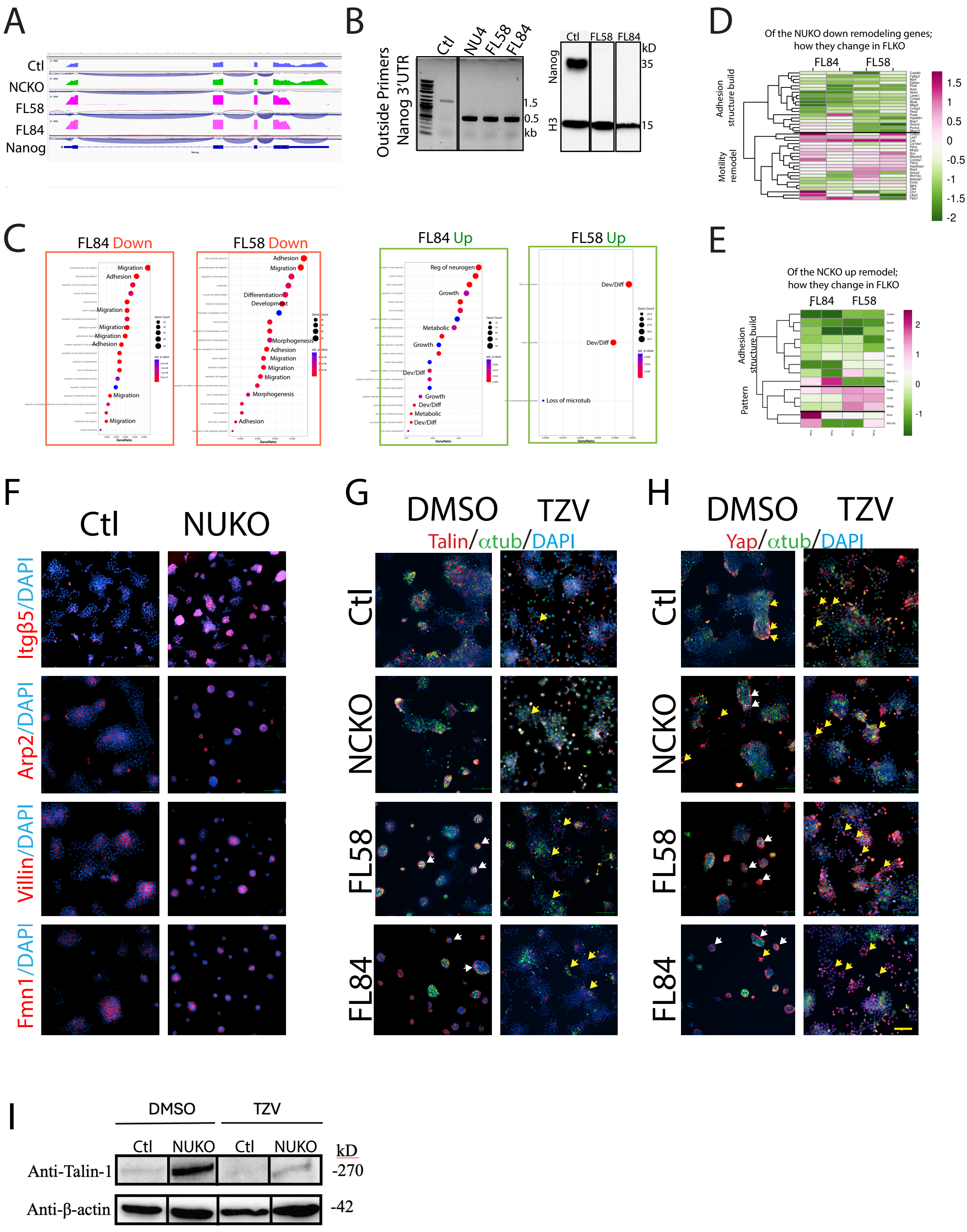


Supp Fig. 5 Yang et al
